## Supplementary material for "Uncertainty quantification in cerebral circulation simulations focusing on the collateral flow: Surrogate model approach with machine learning": S1 Appendix

### S1 Appendix: Details on the design of experiments

The mean and standard deviation (SD) values of the diameters and lengths of the carotid and cerebral arteries of the seven patients (Table 1) were generally similar to those reported in the literature [1-5]. In the case of diameters, we determined that ±50% of the values reported by Liang et al. [6, 7] covered nearly a ±3SD range around the mean of the patients' and literature values; therefore, we adopted this range. The circle of Willis (CoW) is known to exhibit anatomical variations, wherein one or several arteries may be missing [8]. In case of the commonly absent arteries, the lower bound for the diameter was set to 0.1 mm, which allows only a negligible amount of flow ($\bar{Q} \sim 10^{-2}$ mL/min) to pass. By representing the missing artery as an extremely narrow artery with a diameter of 0.1 mm, we could perform simulations efficiently without altering the arterial network topology.

Large differences were observed in the values of arterial lengths between the patient cohort and the literature in the case of the common carotid and vertebral arteries. This can be attributed to racial differences, as most of the values reported in the literature were collected considering Westerners, whereas the seven patients in this study were Asian (Japanese). Therefore, we adopted a range covering the mean ± 3SD of both the patients' and literature values for the common carotid and vertebral arteries. However, for the other arteries, we adopted the ranges of mean ± 3SD of the seven patients only.

The ranges of stenosis parameters considered the mean ±3SD of the seven patients for $D_\mathrm{n}$; 0% (intact) to 100% (occlusion) for $SR$; and 1.0 to 2.699 for $K_\mathrm{t}$ [9]. As indicated in Equation (5), $R_\mathrm{v}$ is maximum when the lumen has the minimum diameter $D_\mathrm{s,min}$ throughout the stenosis.

$$R_\mathrm{v,max} = \frac{128\mu L_\mathrm{s,max}}{\pi D_\mathrm{s,min}^4} = \frac{128\mu L_\mathrm{s,max}}{\pi D_\mathrm{n,min}^4 (1 - SR)^4}, \tag{A1}$$

where $D_\mathrm{n,min}$ denotes the lower bound of $D_\mathrm{n}$, and $L_\mathrm{s,max}$ indicates the maximum stenosis length (assumed to be 40 mm here). To avoid meaningless sampling and ensure that $R_\mathrm{v} \le R_\mathrm{v,max}$, the upper bound of $R_\mathrm{v}$ was defined based on $SR$. Additionally, although $R_\mathrm{v,max}$ approaches infinity as $SR$ approaches 100%, we limited $R_\mathrm{v}$ to less than 500 mmHg s mL$^{-1}$. Even at the maximum possible pressure gradient (approximately 50 mmHg) in vivo, $R_\mathrm{v}$ = 500 mmHg s mL$^{-1}$ resulted in a flow rate of less than 6 mL/min (significantly less than the normal flow rate, which is approximately 257 mL/min [10]); therefore, it represents a nearly occluded lumen. Thus, a higher $R_\mathrm{v}$ has been proven to have no physiological significance.

Although the ranges of the peripheral resistances (PRs) of the CoW were widely set, we ensured that they were physiologically acceptable. For instance, the upper bound of the PR at the middle cerebral artery allows a flow rate of less than 30 mL/min in this artery for pressures defined in the physiological range. Considering that the measured flow rate for the seven patients was 132 ± 27 mL/min, and the flow rate reported in the literature is 146 ± 31 mL/min [10], the range of PR was sufficiently wide to cover a physiologically relevant range of flow rate. Similarly, the range of the scaling factor for the total PR was set such that the simulated mean arterial pressure ranged from 73.3 mmHg (low blood pressure) to 133.3 mmHg (severe hypertension).

[10] Zarrinkoob L, Ambarki K, Wåhlin A, Birgander R, Eklund A, Malm J. Blood flow distribution in cerebral arteries. J Cereb Blood Flow Metab. 2015; 35(4):648–654. https://doi.org/10.1038/jcbfm.2014.241
