## Supplementary material for "Uncertainty quantification in cerebral circulation simulations focusing on the collateral flow: Surrogate model approach with machine learning": S2 Appendix

### S2 Appendix: Details on the uncertainty propagation

As described in the "Uncertainty propagation" subsection in the paper, the probability distribution of $\Delta\bar{Q}_i$ ($i$ = 1, 2, …, 6) under uncertainty was evaluated using the Monte Carlo method, where $\Delta\bar{Q}_i$ indicates the percentage increase of the flow rate at the six outlets of the circle of Willis (CoW) due to the stenosis surgery. We considered the uncertainties in arterial diameters, stenosis parameters, and inflow and outflow measurements of the CoW, which were derived from the clinical data of the patient. In accordance with Fig 4, herein, we denote the uncertain inputs (22 arterial diameters and 8 stenosis parameters) by $\boldsymbol{x}_\mathrm{u} \in \mathbb{R}^{30}$, fixed inputs (22 arterial lengths and age) by $\boldsymbol{x}_\mathrm{f} \in \mathbb{R}^{23}$, and inputs to be iteratively adjusted (6 peripheral resistances of the CoW and scaling factor for the total peripheral resistance) by $\boldsymbol{x}_\mathrm{PR} \in \mathbb{R}^{7}$. Here, $\boldsymbol{x}_\mathrm{u}$, $\boldsymbol{x}_\mathrm{f}$, and $\boldsymbol{x}_\mathrm{PR}$ are column vectors whose concatenation is denoted by $\boldsymbol{x} = [\boldsymbol{x}_\mathrm{u}^\top, \boldsymbol{x}_\mathrm{f}^\top, \boldsymbol{x}_\mathrm{PR}^\top]^\top \in \mathbb{R}^{60}$, where the superscript $\top$ indicates the matrix transpose. Furthermore, the flow and pressure measurements (6 flow rates at the outlets of the CoW and mean arterial pressure), which were used to adjust $\boldsymbol{x}_\mathrm{PR}$, are denoted by $\boldsymbol{y}^\mathrm{target} \in \mathbb{R}^{7}$. The procedure for evaluating the prediction uncertainty in $\Delta\bar{Q}_i$ is described as follows.

#### Step 1: Monte Carlo sampling

The first step in the Monte Carlo method involves the generation of sample values (called "realization") of random variables $\boldsymbol{x}_\mathrm{u}$ and $\boldsymbol{y}^\mathrm{target}$ from the prescribed probability distributions. The sampling of $\boldsymbol{x}_\mathrm{u}^{(s)}$ is straightforward; we generated random real numbers in the intervals shown in Table 2 by assuming a uniform distribution. Here, $\boldsymbol{y}^\mathrm{target}$ includes the target flow rates in the six outlets of the CoW, $\bar{Q}_i^\mathrm{target}$ ($i$ = 1, 2, …, 6), and target mean arterial pressure at the upper arm, $\bar{P}_\mathrm{arm}^\mathrm{target}$. Because the uncertainty associated with $\bar{P}_\mathrm{arm}^\mathrm{target}$ was not considered in this study, $\bar{P}_\mathrm{arm}^{\mathrm{target}(s)}$ is the measured value itself, i.e., $\bar{P}_\mathrm{arm}^{\mathrm{target}(s)} = \bar{P}_\mathrm{arm}^\mathrm{target}$. To sample $\bar{Q}_i^{\mathrm{target}(s)}$, we initially sampled the flow measurements from uncertainty intervals depending on the modality. The flow rates in the six outlets of the CoW, $\bar{Q}_i^{\mathrm{out}(s)}$, were sampled from the uncertainty intervals of single photon emission computed tomography, whereas the flow rates in the three inlets of the CoW, $\bar{Q}_j^{\mathrm{in}(s)}$ ($j$ = 1, 2, 3), were sampled from the uncertainty intervals of phase contrast magnetic resonance imaging or ultrasound measurements. Subsequently, $\bar{Q}_i^{\mathrm{target}(s)}$ was calculated in accordance with Equation (8):

$$\bar{Q}_i^{\mathrm{target}(s)} = \bar{Q}_i^{\mathrm{out}(s)} \cdot \frac{\sum_{j=1}^{3} \bar{Q}_j^{\mathrm{in}(s)}}{\sum_{k=1}^{6} \bar{Q}_k^{\mathrm{out}(s)}}. \tag{A2}$$

Thus, $\boldsymbol{y}^{\mathrm{target}(s)}$ was sampled as $[\bar{Q}_1^{\mathrm{target}(s)}, \bar{Q}_2^{\mathrm{target}(s)}, \ldots, \bar{Q}_6^{\mathrm{target}(s)}, \bar{P}_\mathrm{arm}^{\mathrm{target}(s)}]^\top$.

#### Step 2: Preoperative adjustment

The surrogate model predicts outputs $\boldsymbol{y}^{(s)} \in \mathbb{R}^{45}$ based on the given inputs $\boldsymbol{x}^{(s)} = [\boldsymbol{x}_\mathrm{u}^{(s)\top}, \boldsymbol{x}_\mathrm{f}^\top, \boldsymbol{x}_\mathrm{PR}^{(s)\top}]^\top \in \mathbb{R}^{60}$. The outputs include the flow rates at the six outlets of the CoW, $\bar{Q}_i^{(s)}$, and the mean arterial pressure, $\bar{P}_\mathrm{arm}^{(s)}$. The purpose of this step is to evaluate the values of $\boldsymbol{x}_\mathrm{PR}^{*(s)} = [PR_1^{*(s)}, PR_2^{*(s)}, \ldots, PR_6^{*(s)}, \alpha_\mathrm{PR}^{*(s)}]^\top$ that minimize the difference between $\bar{Q}_i^{(s)}$ and $\bar{Q}_i^{\mathrm{target}(s)}$ and between $\bar{P}_\mathrm{arm}^{(s)}$ and $\bar{P}_\mathrm{arm}^{\mathrm{target}(s)}$. Herein, $PR_i^{(s)}$ ($i$ = 1, 2, …, 6) are the peripheral resistances (PRs) at the six outlets of the CoW, and $\alpha_\mathrm{PR}^{(s)}$ is the scaling factor for the total PR. To obtain $\boldsymbol{x}_\mathrm{PR}^{*(s)}$, we iteratively adjusted $\boldsymbol{x}_\mathrm{PR}^{(s)}$ based on the difference between the predicted and target quantities of the flow rate and pressure [1, 2]:

$$[PR_i^{(s)}]^{n+1} = [PR_i^{(s)}]^{n} \left( 1 + c \cdot \frac{[\bar{Q}_i^{(s)}]^{n} - \bar{Q}_i^{\mathrm{target}(s)}}{\bar{Q}_i^{\mathrm{target}(s)}} \right), \tag{A3}$$

$$[\alpha_\mathrm{PR}^{(s)}]^{n+1} = [\alpha_\mathrm{PR}^{(s)}]^{n} \left( 1 - c \cdot \frac{[\bar{P}_\mathrm{arm}^{(s)}]^{n} - \bar{P}_\mathrm{arm}^{\mathrm{target}(s)}}{\bar{P}_\mathrm{arm}^{\mathrm{target}(s)}} \right), \tag{A4}$$

where $[\cdot]^n$ denotes the values at the $n$-th iteration, and $c$ is the relaxation coefficient (taken to be 0.9). The initial values for $PR_i^{(s)}$ were considered as the values reported in the literature [3], and the initial value for $\alpha_{\mathrm{PR}}^{(s)}$ was set to 1. The iterations were continued until the following convergence criteria were met:

$$\left|\frac{[\bar{Q}_i^{(s)}]^n - \bar{Q}_i^{\mathrm{target}(s)}}{\bar{Q}_i^{\mathrm{target}(s)}}\right| < \varepsilon, \tag{A5}$$

$$\left|\frac{[\bar{P}_{\mathrm{arm}}^{(s)}]^n - \bar{P}_{\mathrm{arm}}^{\mathrm{target}(s)}}{\bar{P}_{\mathrm{arm}}^{\mathrm{target}(s)}}\right| < \varepsilon, \tag{A6}$$

where $\varepsilon$ is the tolerance error, which was set to 0.005 in this study. The converged solution of $\boldsymbol{x}_{\mathrm{PR}}^{*(s)}$ was regarded as the patient's preoperative PRs, and the preoperative flow rates and pressures, $\boldsymbol{y}^{\mathrm{pre}(s)}$, were obtained using the inputs $\boldsymbol{x}^{(s)} = [\boldsymbol{x}_{\mathrm{u}}^{(s)\top}, \boldsymbol{x}_{\mathrm{f}}^{\top}, \boldsymbol{x}_{\mathrm{PR}}^{*(s)\top}]^{\top}$.

Note that Equations (A5) and (A6) may not be satisfied for certain samples, as we consider a wide uncertainty interval. For example, the combination of $\boldsymbol{x}_{\mathrm{u}}^{(s)}$, representing extremely severe stenosis, and $\boldsymbol{y}^{\mathrm{target}(s)}$, indicating a much larger flow rate in the stenosis side than the normal side, is physically unrealizable. Therefore, we set the maximum number of iterations as $n = 400$; samples that did not satisfy the criteria within these iterations were rejected.

#### Step 3: Postoperative prediction

Following parameter adjustment for reproducing the patient's preoperative cerebral circulation, this step simulated the surgical dilation of the stenosis and predicted the cerebral circulation immediately after the surgery. All inputs for the prediction are the same as those in the previous step, except the stenosis parameters in $\boldsymbol{x}_{\mathrm{u}}^{(s)}$, which were modified to $R_{\mathrm{v}} = 0$, $D_{\mathrm{n}} = D_{\mathrm{ICA}}$ (diameter of the internal carotid artery), $SR = 0$, and $K_{\mathrm{t}} = 0$ to reflect the complete dilation of the stenosis. As depicted in Fig 4, the inputs used for postoperative prediction can be written as $\boldsymbol{x}^{(s)} = [\boldsymbol{x}_{\mathrm{u}}^{*(s)\top}, \boldsymbol{x}_{\mathrm{f}}^{\top}, \boldsymbol{x}_{\mathrm{PR}}^{*(s)\top}]^{\top}$, where $\boldsymbol{x}_{\mathrm{u}}^{*(s)}$ denotes the sampled inputs with modified stenosis parameters. According to these inputs, the surrogate model predicted the patient's postoperative flow rates and pressures, $\boldsymbol{y}^{\mathrm{post}(s)}$.

In the postoperative prediction, it was assumed that the surgery did not alter the arterial geometry (except for stenosis) and PRs. This assumption is justified because we aim to predict the cerebral circulation "immediately after" the surgery. Additionally, autoregulation and remodeling of the cerebral arteries generally prevent an abrupt change in blood flow. Therefore, our assumption is appropriate for predicting the maximum possible $\Delta\bar{Q}_i$, which is the most dangerous surgical outcome in terms of cerebral hyperperfusion.

#### Step 4: Statistics evaluation and sample addition

Finally, $\Delta\bar{Q}_i^{(s)}$ was calculated based on $\boldsymbol{y}^{\mathrm{pre}(s)}$ and $\boldsymbol{y}^{\mathrm{post}(s)}$, as follows:

$$\Delta\bar{Q}_i^{(s)} = \frac{\bar{Q}_i^{\mathrm{post}(s)} - \bar{Q}_i^{\mathrm{pre}(s)}}{\bar{Q}_i^{\mathrm{pre}(s)}} \times 100\%, \quad i = 1, 2, \ldots, 6, \tag{A7}$$

where $\bar{Q}_i^{\mathrm{pre}(s)}$ and $\bar{Q}_i^{\mathrm{post}(s)}$ are the flow rates in the six outlets of the CoW in $\boldsymbol{y}^{\mathrm{pre}(s)}$ and $\boldsymbol{y}^{\mathrm{post}(s)}$, respectively. By repeating Step 1 through Step 3 for $N_{\mathrm{MC}}$ times, the Monte Carlo samples of $\Delta\bar{Q}_i^{(s)}$ ($s = 1, 2, \ldots, N_{\mathrm{MC}}$) were obtained. The statistics of $\Delta\bar{Q}_i$ under uncertainties were estimated using the collected samples. For example, the mean of $\Delta\bar{Q}_i$ was calculated as follows:

$$\mathbb{E}[\Delta\bar{Q}_i] \approx \mathbb{E}_{\mathrm{MC}}[\Delta\bar{Q}_i] = \frac{1}{N_{\mathrm{MC}}}\sum_{s=1}^{N_{\mathrm{MC}}} \Delta\bar{Q}_i^{(s)}, \tag{A8}$$

and the variance of $\Delta\bar{Q}_i$ was calculated as follows:

$$\mathbb{V}[\Delta\bar{Q}_i] \approx \mathbb{V}_{\mathrm{MC}}[\Delta\bar{Q}_i] = \frac{1}{N_{\mathrm{MC}}}\sum_{s=1}^{N_{\mathrm{MC}}}\left(\Delta\bar{Q}_i^{(s)} - \mathbb{E}_{\mathrm{MC}}[\Delta\bar{Q}_i]\right). \tag{A9}$$

Furthermore, the probability distribution of $\Delta\bar{Q}_i$ can be written as follows:

$$\rho(\Delta\bar{Q}_i) \approx \rho_{\mathrm{MC}}(\Delta\bar{Q}_i) = \frac{1}{N_{\mathrm{MC}}}\sum_{s=1}^{N_{\mathrm{MC}}}\delta\left(\Delta\bar{Q}_i - \Delta\bar{Q}_i^{(s)}\right), \tag{A10}$$

where $\delta$ is the Dirac delta function. Equation (A10) corresponds to a histogram, which depicts the frequencies of $\Delta\bar{Q}_i^{(s)}$ normalized by the total sample size.

The number of Monte Carlo samples, $N_{\mathrm{MC}}$, was increased sequentially until the statistics of $\Delta\bar{Q}_i$ converged. As a basic policy, we increased $N_{\mathrm{MC}}$ by 10 000 and ensured that the change in mean and variance of $\Delta\bar{Q}_i$ was within 0.1%. We also confirmed that there was no significant change in the probability of $\Delta\bar{Q}_i > 100\%$ when $N_{\mathrm{MC}}$ was increased.

#### References


[1] Liang F, Oshima M, Huang H, Liu H, Takagi S. Numerical study of cerebroarterial hemodynamic changes following carotid artery operation: a comparison between multiscale modeling and stand-alone three-dimensional modeling. J Biomech Eng. 2015; 137(10):101011. https://doi.org/10.1115/1.4031457

[2] Zhang H, Fujiwara N, Kobayashi M, Yamada S, Liang F, Takagi S, et al. Development of a numerical method for patient-specific cerebral circulation using 1D–0D simulation of the entire cardiovascular system with SPECT data. Ann Biomed Eng. 2016; 44(8):2351–2363. https://doi.org/10.1007/s10439-015-1544-8

[3] Liang F, Fukasaku K, Liu H, Takagi S. A computational model study of the influence of the anatomy of the circle of Willis on cerebral hyperperfusion following carotid artery surgery. Biomed Eng Online. 2011; 10:84. https://doi.org/10.1186/1475-925X-10-84
