## Supplementary figures and images for "Uncertainty quantification in cerebral circulation simulations focusing on the collateral flow: Surrogate model approach with machine learning"

### S1 Fig

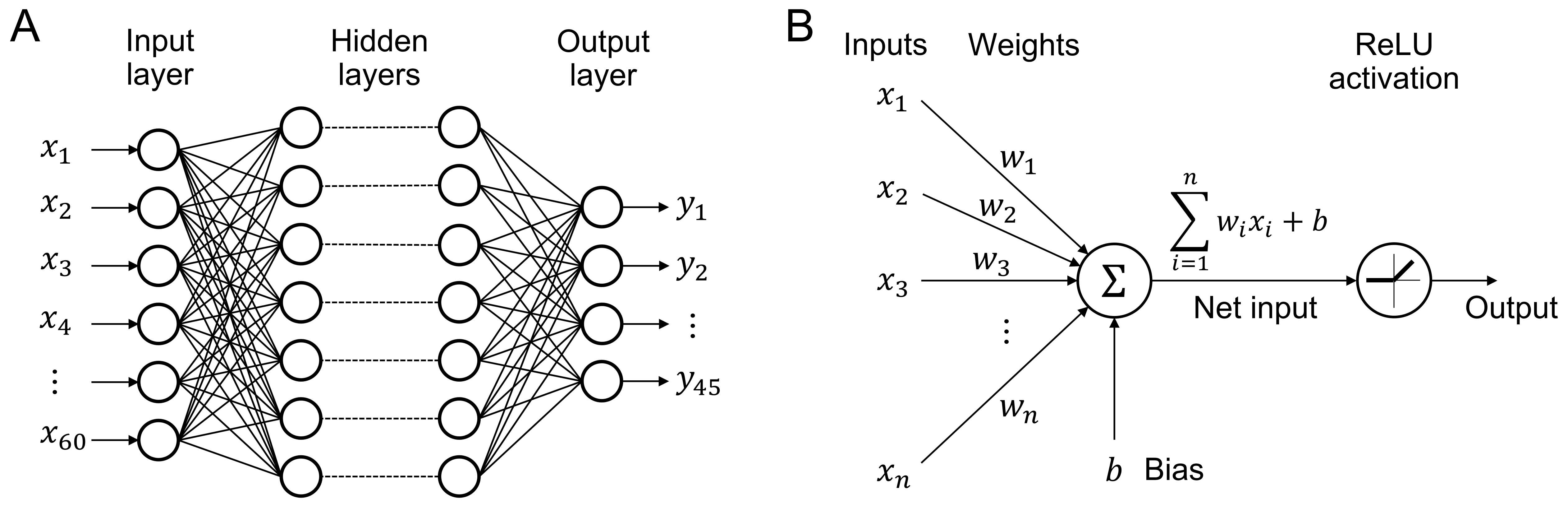

### S2 Fig

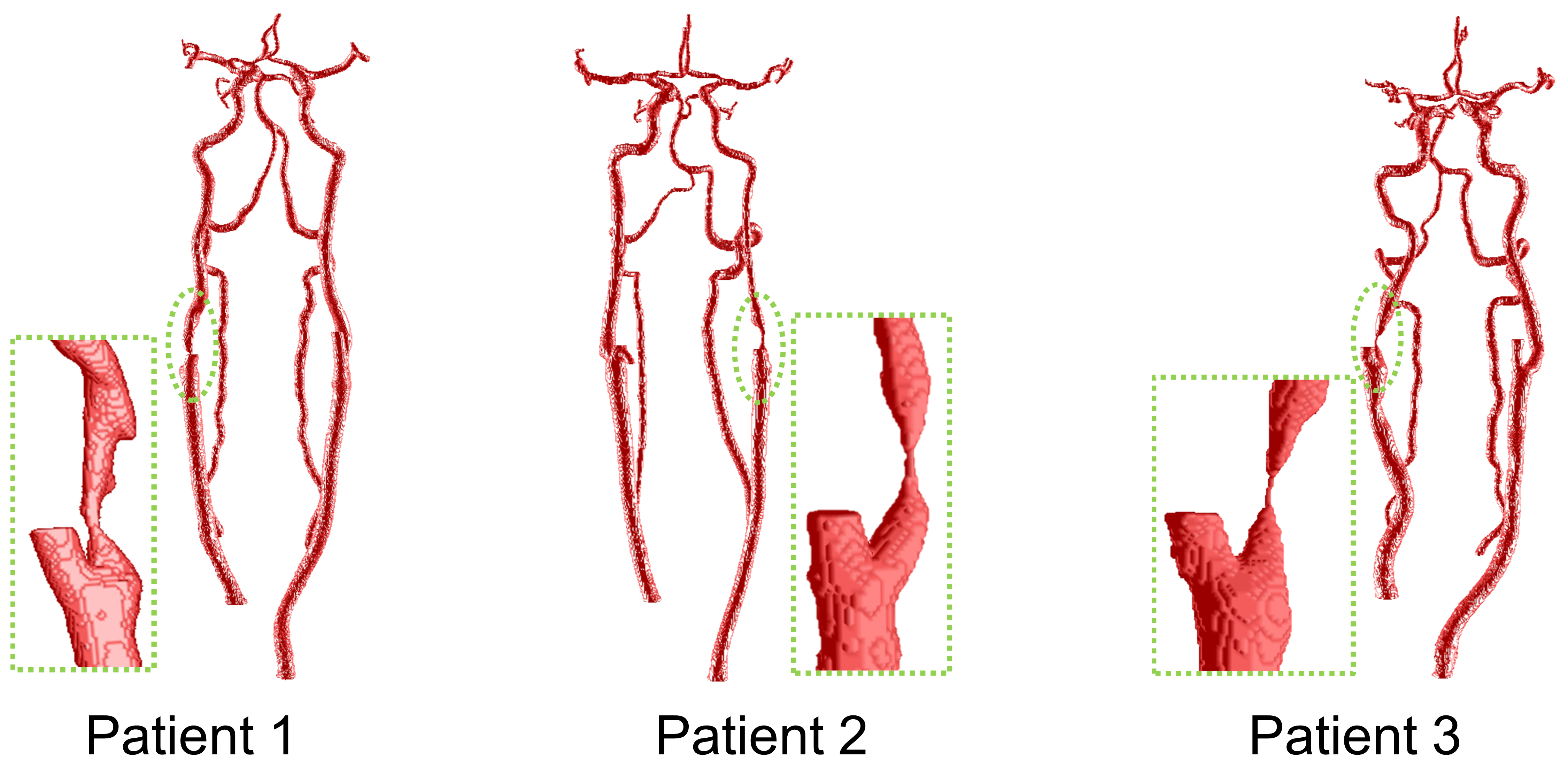

### S3 Fig

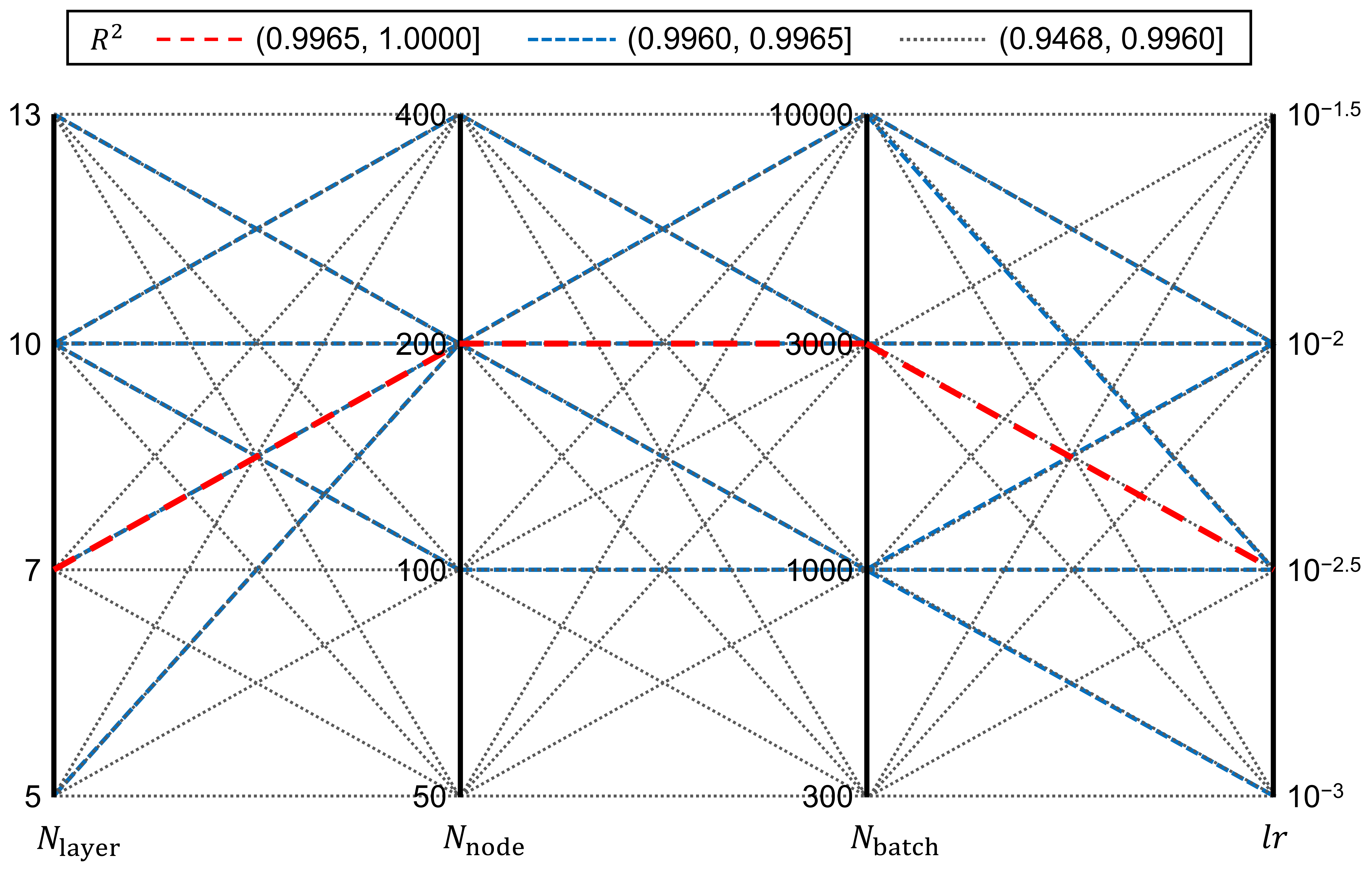

### S4 Fig

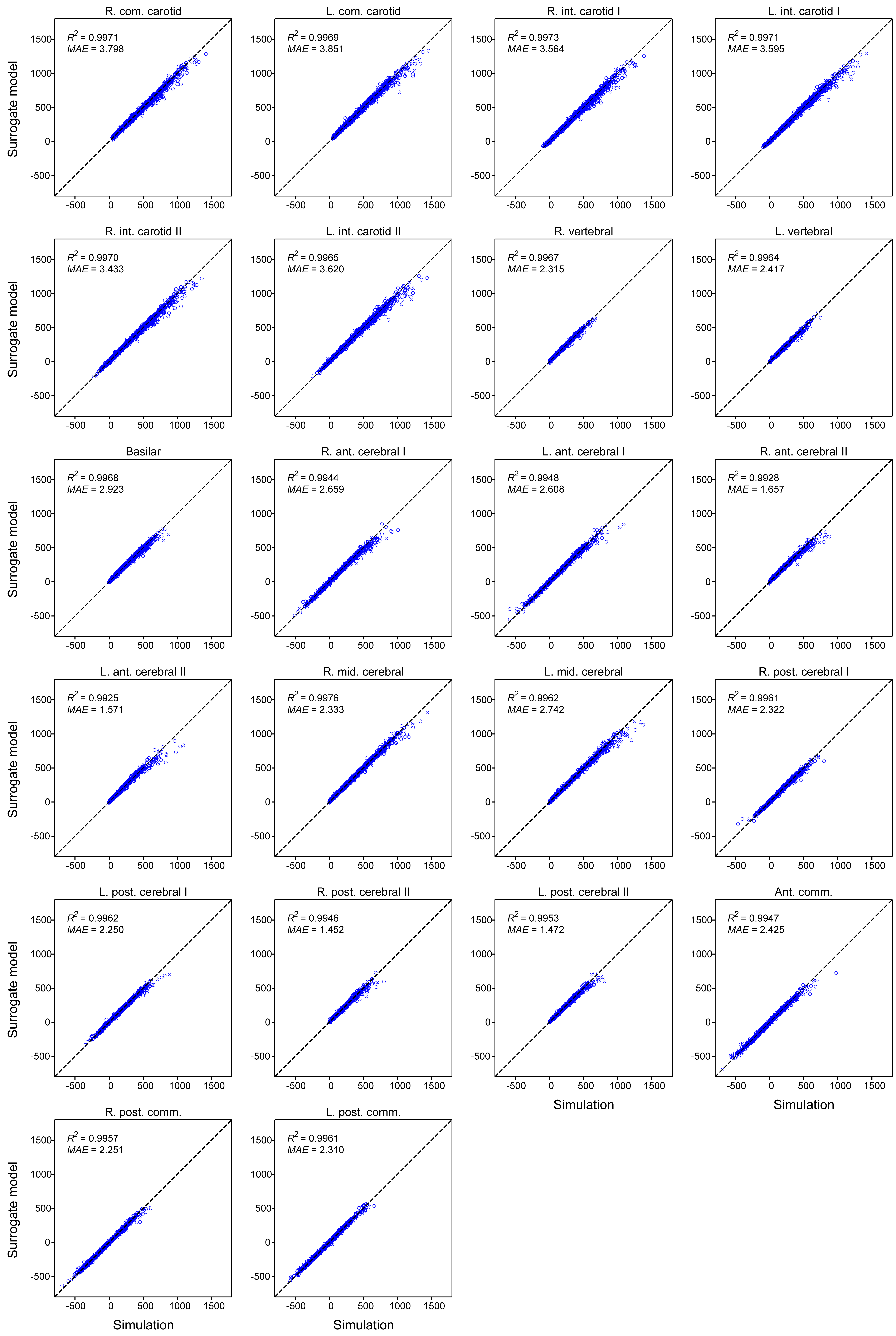

### S5 Fig

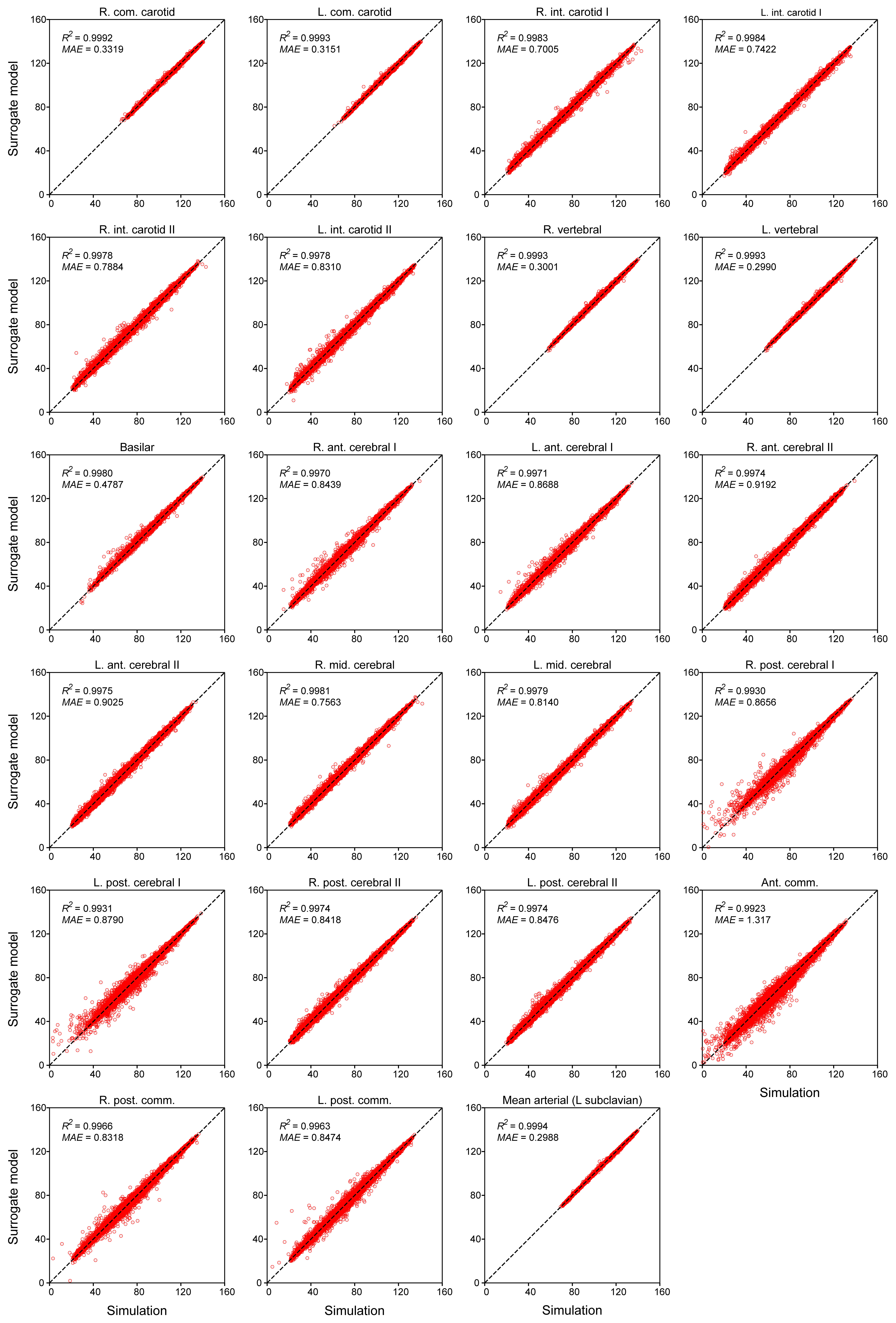
